# High-precision CRISPR-Cas9 base editors with minimized bystander and off-target mutations

**DOI:** 10.1101/273938

**Authors:** Jason M. Gehrke, Oliver Cervantes, M. Kendell Clement, Luca Pinello, J. Keith Joung

## Abstract

Recently described base editor (**BE**) technology, which uses CRISPR-Cas9 to direct cytidine deaminase enzymatic activity to specific genomic loci, enables the highly efficient introduction of precise cytidine-to-thymidine (C → T) DNA alterations in many different cell types and organisms^1–6^. In contrast to genome-editing nucleases^7–9^, BEs avoid the need to introduce double-strand breaks or exogenous donor DNA templates and induce lower levels of unwanted variable-length insertion/deletion mutations (indels)^1^,^2^,^10^. However, existing BEs can also efficiently create unwanted C to T alterations when more than one C is present within the five base pair “editing window” of these proteins, a lack of precision that can cause potentially deleterious bystander mutations. Mutations in the cytidine deaminase enzyme can shorten the length of the editing window and thereby partially address this limitation but these BE variants still do not discriminate among multiple cytidines within the narrowed window and also possess a more limited targeting range^11^. Here, we describe an alternative strategy for reducing bystander mutations using a novel BE architecture that harbors an engineered human APOBEC3A (**eA3A**) domain, which preferentially deaminates cytidines according to a TCR>TCY>VCN (V = G, A, C, Y = C, T) hierarchy. In direct comparisons with the widely used BE3 fusion in human cells, our eA3A-BE3 fusion exhibits comparable activities on cytidines in TC motifs but greatly reduced or no significant editing on cytidines in other sequence contexts. Importantly, we show that eA3A-BE3 can correct a human beta-thalassemia promoter mutation with much higher (>40-fold) precision than BE3, substantially minimizing the creation of an undesirable bystander mutation. Surprisingly, we also found that eA3A-BE3 shows reduced mutation frequencies on known off-target sites of BE3, even when targeting promiscuous homopolymeric sites. Our results validate a general strategy to improve the precision of base editors by engineering their cytidine deaminases to possess greater sequence specificity, an important proof-of-principle that should motivate the development of a larger suite of new base editors with such properties.

To engineer base editor fusions with greater precision within the editing window, we sought to leverage the natural diversity of cytidine deaminase proteins to employ a deaminase with greater sequence specificity than the rat APOBEC1 (**rAPO1**) deaminase present in the widely used BE3 architecture. BE3 consists of a *Streptococcus pyogenes* Cas9 nuclease bearing a mutation that converts it into a nickase (**nCas9**) fused to rAPO1 and a uracil glycosylase inhibitor (**UGI**) (**Fig. 1a**). We replaced the rAPO1 in BE3 with the human APOBEC3A (A3A) cytidine deaminase to create A3A-BE3 (**Fig. 1a**). We used A3A because previously published *in vitro* biochemical studies showed that it preferentially deaminates cytidines embedded in the context of a TCR trinucleotide motif (where R = A/G)^12–14^. To test the precision of base editing by A3A-BE3, we used a guide RNA targeted to a site in a single integrated *EGFP* reporter gene in human U2OS cells that bears both a cognate motif (TCG) and a non-cognate bystander (GCT) motif within the expected editing window. Surprisingly, A3A-BE3 did not exhibit preferential base editing of the cytidine in the TCG motif over the GCT motif (**Fig. 1b**).

**Figure 1:**
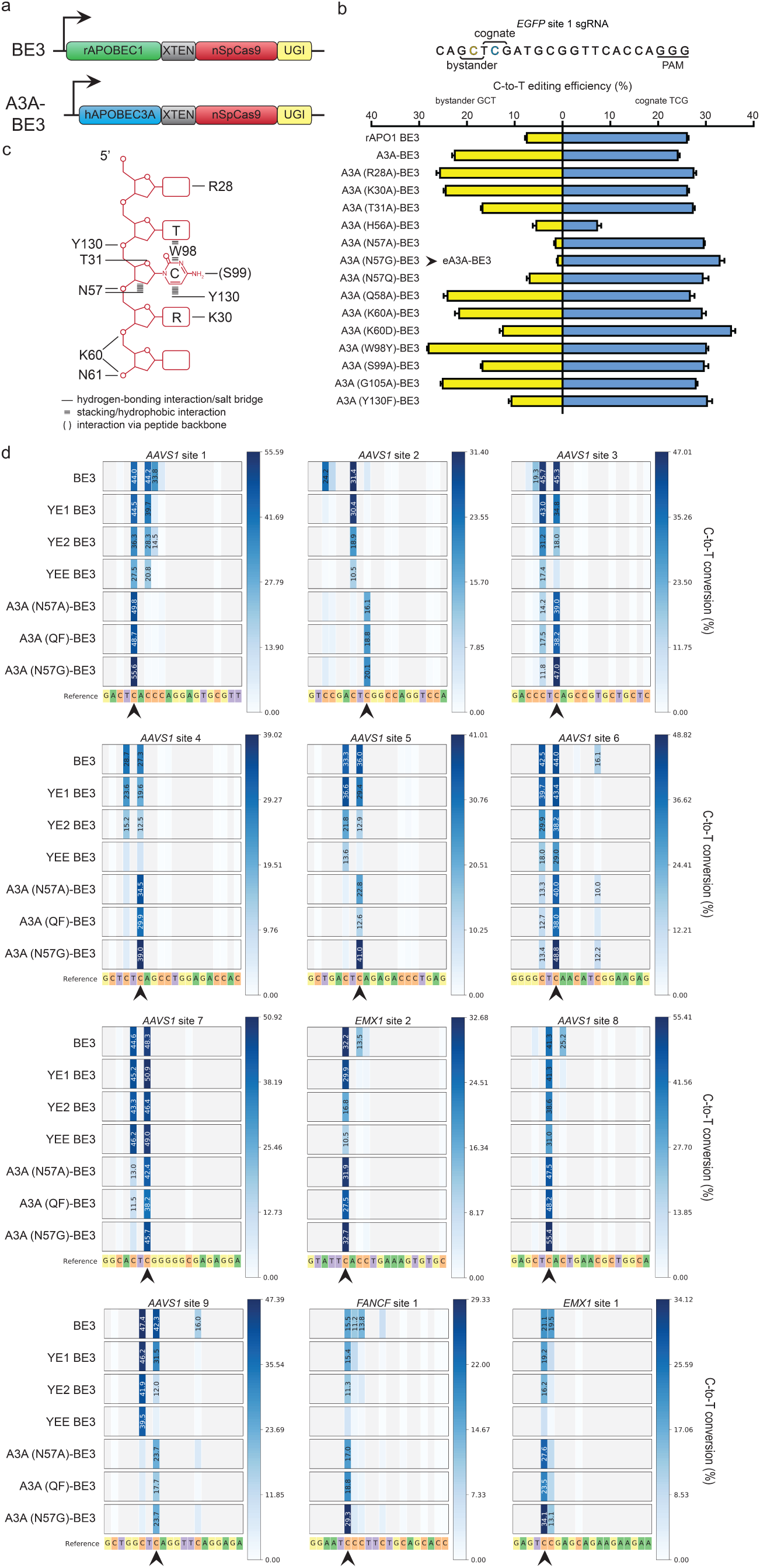
Engineering and characterization of an A3A-BE3 base editor that selectively edits Cs preceded by a 5’ T. (a) Schematic illustrating the architecture of the original BE3 fusion (consisting of rAPO1 linked to SpCas9 nickase and UGI) and the A3A-BE3 fusion. (b) Activities of BE3, A3A-BE3, and a series of A3A-BE3 variants bearing mutations in A3A on an integrated *EGFP* reporter gene target site bearing a cognate cytidine preceded by a 5’ T and a bystander cytidine preceded by a 5‘ G in the editing window. (c) Schematic summarizing non-specific interactions between amino acid positions in A3A and its substrate single-stranded DNA derived from previously published co-crystal structures. (d) Heat maps showing C-to-T editing efficiencies for BE3, YE BE3s, and various A3A-BE3 variants at 12 endogenous human gene target sites, each bearing a cognate cytidine preceded by a 5’ T (indicated with a black arrow) and one or more bystander cytidines within the editing window. Editing efficiencies shown represent the mean of three biological replicates.

We hypothesized that the apparent loss of sequence preference by A3A on the *EGFP* site in the context of base editor fusion might have been due to its increased proximity secondary to recruitment to that locus. We envisioned that sequence selectivity might be restored by reducing the non-specific binding energy of A3A for its substrate DNA. Based on co-crystal structures of A3A and a single-stranded DNA substrate, we identified 11 residues in the deaminase that appear to mediate non-specific contacts to the DNA (**Fig. 1c**)^13^,^14^ and created a series of 14 mutant A3A proteins bearing one or more amino acid substitutions at these positions (**Fig. 1b**). Testing of these mutated A3A proteins in the context of the A3A-BE3 architecture showed that most retained high activity on both the bystander and cognate motifs but that those bearing mutations in position N57 drastically reduced bystander motif alteration while retaining near-wild-type activity on the cognate motif (**Fig. 1b**). We also found we could further increase the preference of A3A for its cognate TCR motif by combining point mutations at residues N57, K60, or Y130 (**Supplementary Fig. 1**). This strategy yielded the N57Q/Y130F (QF) variant, which appeared to have similar sequence preferences to the N57A/G single mutation variants. We conclude that engineering mutations into A3A enables restoration of its cytidine deaminase sequence preference in the context of a BE fusion.

We next sought to more broadly test the precision of our A3A-BE3 fusions on a larger number of sites within endogenous human genes. To do this, we used 12 different gRNAs targeted to three different human genes and directly compared the editing activities of seven base editor fusions: three A3A-BE3 variants (bearing N57G, N57A, and N57Q/Y130F mutations in A3A), the original BE3, and three previously described BE3 variants, YE1, YE2, and YEE BE3 (YE BE3s), that possess point mutations in rAPO1 designed to slow its kinetic rate and thereby restrict the editing window^11^ (**Fig. 1d**). We found that among all seven base editor fusions tested, A3A (N57G)-BE3 displayed the highest activity at cognate motifs while minimizing bystander cytidine editing at all of the sites tested. At eight of 12 tested sites, A3A (N57G)-BE3 induced 5- to 264-fold (median of 12.8-fold) editing of cognate motifs compared to bystander motifs in the editing window. At the other four sites, A3A (N57G)-BE3 induced much lower frequencies of editing at bystander motifs than that observed with BE3 while retaining high activity at the cognate motif. As expected, all three A3A-BE3 variants maintained a five-nucleotide editing window similar to that of the wild-type A3A-BE3 enzyme. YE1 BE3 narrowed the editing window to approximately three nucleotides in most cases while still retaining catalytic activity at the cognate motif similar to BE3. YE2 BE3 failed to produce fewer bystander mutations compared to YE1 BE3, and YEE BE3 lost significant activity at 9 sites. Based on these results, we chose the A3A (N57G)-BE3 variant for additional characterization and refer to it hereafter as **eA3A-BE3** (for engineered A3A-BE3).

To examine the purity of the edited alleles produced by eA3A-BE3, we performed a detailed analysis of the high-throughput sequencing results obtained at the 12 endogenous human gene target sites (**Fig. 1d**), which revealed that eA3A-BE3 induced significant differences in the frequencies of unwanted alterations compared with the original BE3. At 11 of the 12 sites, eA3A-BE3 showed an altered frequency of unwanted base substitutions (i.e., C to A or G) from what was observed with the original BE3 fusion (**Supplementary Fig. 2a**). This finding provides additional support for the previously proposed hypothesis that processing of genomic lesions with multiple uracils by endogenous DNA repair machinery differs from those with single uracils^10^. In addition, we observed eA3A-BE3 also induced fewer indels than BE3 at eight of the twelve tested sites (**Supplementary Fig. 2b**), suggesting that single nucleotide editing does not generally produce indels at substantially different frequencies of indels than multi-base editing.

To attempt to improve the precision of eA3A-BE3 at sites that had cognate-to-bystander editing ratios of less than five, we sought to further reduce the catalytic efficiency of the A3A N57G deaminase.Our rationale for this strategy stemmed from the observation that at these four sites the majority of bystander deamination appears to co-occur with cognate deamination: bystander deamination without cognate deamination is found at fewer than 2% of all alleles at these sites while deamination of the cognate cytidine alone was found in at least 20% of alleles (**Supplementary Figure 3**). To obtain a protein with lower catalytic rate, we added to eA3A mutations to the homologous positions for three previously described residues (E38, A71 or I96) that modulate the catalytic activity of the human AID enzyme^15^. We then screened these mutated eA3A-BE3 variants for activity on three of the genomic sites that had retained significant bystander deamination when edited with eA3A-BE3 (**Supplementary Figure 4**). Mutations made to residues I96 and A71 greatly decreased mutation of bystander motifs at each of the three target sites while retaining 50-75% of A3A N57G BE3 activity at the cognate motif. These results suggest that it may be possible to further modify eA3A-BE3 using a set of defined mutations to tune precision at sites with less than optimal cognate-to-bystander editing ratios.

Having characterized the on-target activity of eA3A-BE3, we next sought to characterize and optimize its potential off-target activity. To do this, we used three different gRNAs (targeted to the *EMX1, FANCF*, and *VEGFA* genes) (**Supplementary Table 1**), for which a total of 60 off-targets had been previously identified with the original BE3 by either Digenome-seq (performed with rAPO1-nSpCas9 also known as “BE3deltaUGI”^16^ or GUIDE-seq (performed with SpCas9 nuclease)^1^,^17^. Using GUIDE-seq performed with SpCas9 nuclease, we also identified two potential off-target sites for a a fourth gRNA (targeted to the *CTNNB1* gene) (**Supplementary Fig. 5**). Finally, we identified some additional closely matched sequences for the *CTNNB1*-targeting gRNA *in silico* in the human reference genome using the Cas-OFFinder program^18^. We then performed targeted amplicon sequencing of these 60 sites to assess base editing events induced by the original BE3 and eA3A-BE3 with the four gRNAs in human HEK293T cells. For all four gRNAs, on-target base editing efficiency with eA3A-BE3 either matched or outperformed the original BE3 (**Figs. 2a - 2d**). For 36 of the 60 potential off-target sites we examined, BE3 induced significant base editing events (compared to control amplicons from untransfected cells) (**Figs. 2a – 2d** and **Supplementary Table 2**). Surprisingly, at 34 of these 36 off-target sites, eA3A-BE3 induced significantly lower frequencies of base editing events with no significant detectable editing at 21 of these 36 sites (**Figs. 2a – 2d**). The N57G mutation in the A3A deaminase part of eA3A-BE3 is critical for this higher specificity because the A3A-BE3 fusion lacking this alteration showed higher off-target mutations with the *EMX1* site 1 and *FANCF* site 1 gRNAs (**Supplementary Figure 6**. The addition of mutations previously shown to improve the genome-wide specificity of SpCas9 (the “HF1” and “Hypa” mutations^19^,^20^) together with an additional second UGI domain further reduced the off-target base editing events (reducing them to undetectable levels for all but 5 of the 15 sites that still showed detectable edits with eA3A-BE3) (**Figs. 2a – 2d**). These higher-specificity variants also induce improved base editing product purity and reduced frequencies of indels at on-target sites (**Supplementary Figure 7**), consistent with earlier studies that used similar strategies to achieve these outcomes for the original BE3 fusion^10^,^21^.

**Figure 2:**
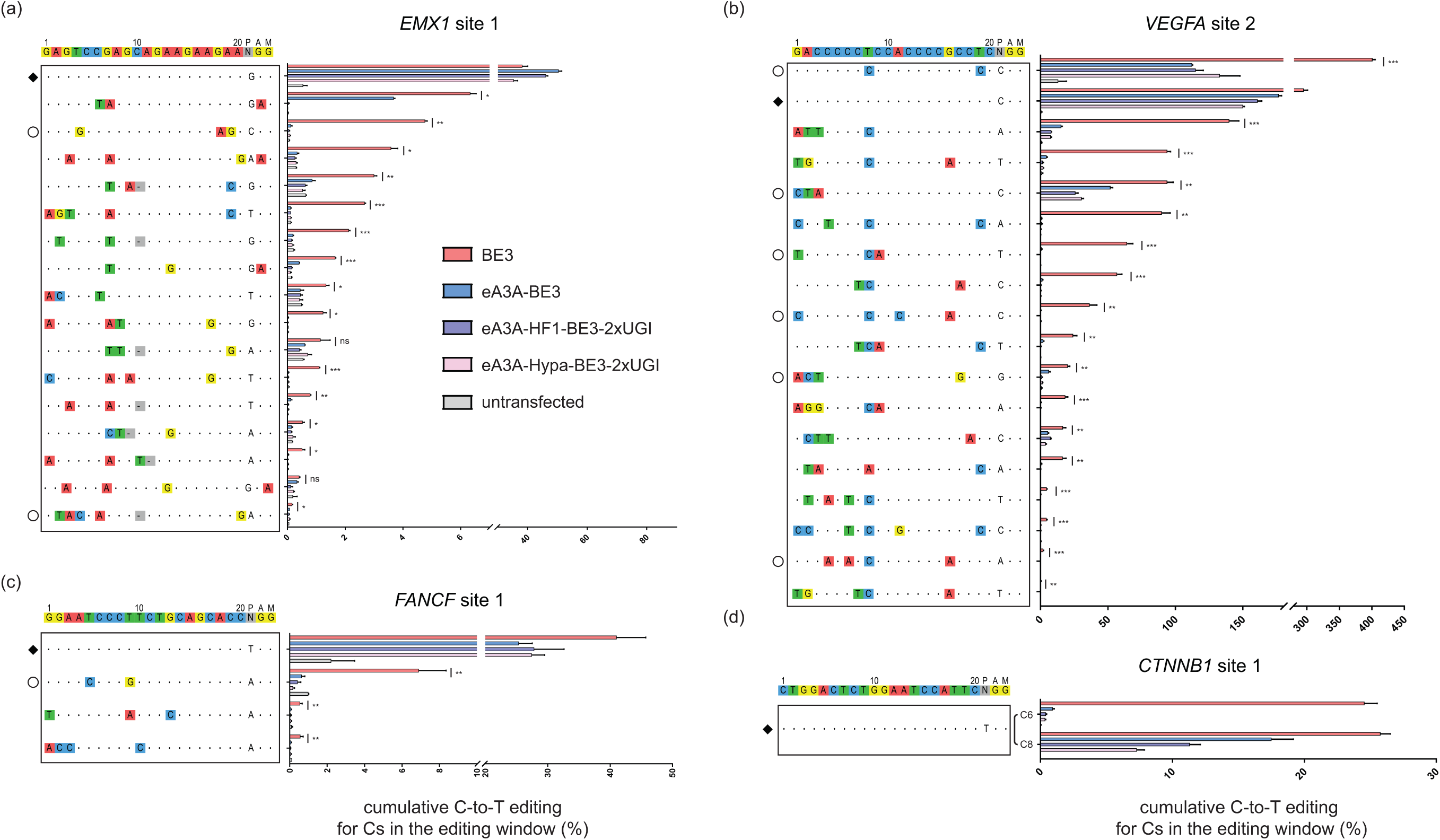
Off-target editing activities of BE3 and eA3A-BE3 variants. On- and off-target editing frequencies of four gRNAs targeted to (a) *EMX1* site 1, (b) *VEGFA* site 2, (c) *FANCF* site 1 or (d) *CTNNB1* site 1 with BE3 or one of the indicated eA3A-BE3 variants. Percentage edits represent the sum of all edited Cs in the editing window and represent the mean of three biological replicates with error bars representing SEMs. Intended target sequence is shown at the top of each graph. On-target sites are marked with a black diamond to the left and mismatches or bulges in the various off-target sites are shown with colored boxes or a dash in gray boxes, respectively. Off-target sites that lose the cognate TC motif within the editing window and thus might be expected to show lower off-target editing by eA3A, are noted with empty circles to the left. Asterisks indicate statistically significant differences in editing efficiencies observed between BE3 and eA3A-BE3 at each site (* p < 0.05, ** p < 0.005, *** p < 0.0005).

To test the eA3A-BE3 fusion on a disease-relevant mutation, we examined its activity on a common β-thalassemia allele found in China and some Southeast Asian populations^22^,^23^ for which single nucleotide editing is critical. Mutation of position -28 of the human *HBB* promoter from an A to G (and therefore a T to C on the complementary strand) results in β-thalassemia disease (**Fig. 3a**). The *HBB* -28 C mutation can be corrected with SpCas9 and a gRNA with the C falling within the predicted editing window. However, another C (at position -25 of the *HBB* promoter) is also present within the editing window of this gRNA and previous work has shown that mutation of this base can cause a β-thalassemia phenotype in humans independent of the nucleotide identity at the -28 position (**Fig. 3a**)^24^,^25^.We directly compared the abilities of the original BE3, the YE BE3s, and eA3A-BE3 to edit an integrated copy of 200 bps of mutant HBB promoter sequence encompassing the cytidines at positions -28 and -25 in HEK293T cells. (For technical reasons, all experiments targeting the *HBB* -28 (A>G) allele used a gRNA expressed with a self-cleaving hammerhead ribozyme on its 5‘ end (**Online Methods**).) As expected, eA3A-BE3 showed higher precision than BE3 and the YE BE3s for selectively editing the -28 cytidine relative to the -25 cytidine (**Fig. 3b**). This resulted in substantially higher levels of perfectly corrected alleles bearing only a -28 C to T edit: 22.48% for eA3A-BE3 compared with 0.57%, 1.04%, 0.92%, and 0.76% for BE3, YE1 BE3, YE2 BE3, and YEE BE3, respectively (**Fig. 3c**). Analysis of eight potential off-target sites for the *HBB*-targeted gRNA (three identified by GUIDE-seq with SpCas9 nuclease (**Supplementary Fig. 5** and five by *in silico* methods; **Online Methods**) showed that eA3A-BE3 induced significant off-target editing at two sites while BE3 induced significant editing at these same two sites and an additional third site all at higher frequencies (**Fig. 3d**). As expected, use of the eA3A-HF1-BE3-2xUGI and eA3A-Hypa-BE3-2xUGI fusions with the *HBB* gRNA reduced the frequencies of off-target edits to undetectable levels at all eight sites we examined (**Fig. 3d**). Both high-fidelity base editor fusions also improved product purity, resulting in a reduction of unwanted -28 C to G edits that are also known to cause β-thalassemia from 16.3% with eA3A-BE3 to 8.8% and 7.5% with the HF1 and Hypa variants, respectively (**Fig. 3c**).

**Figure 3:**
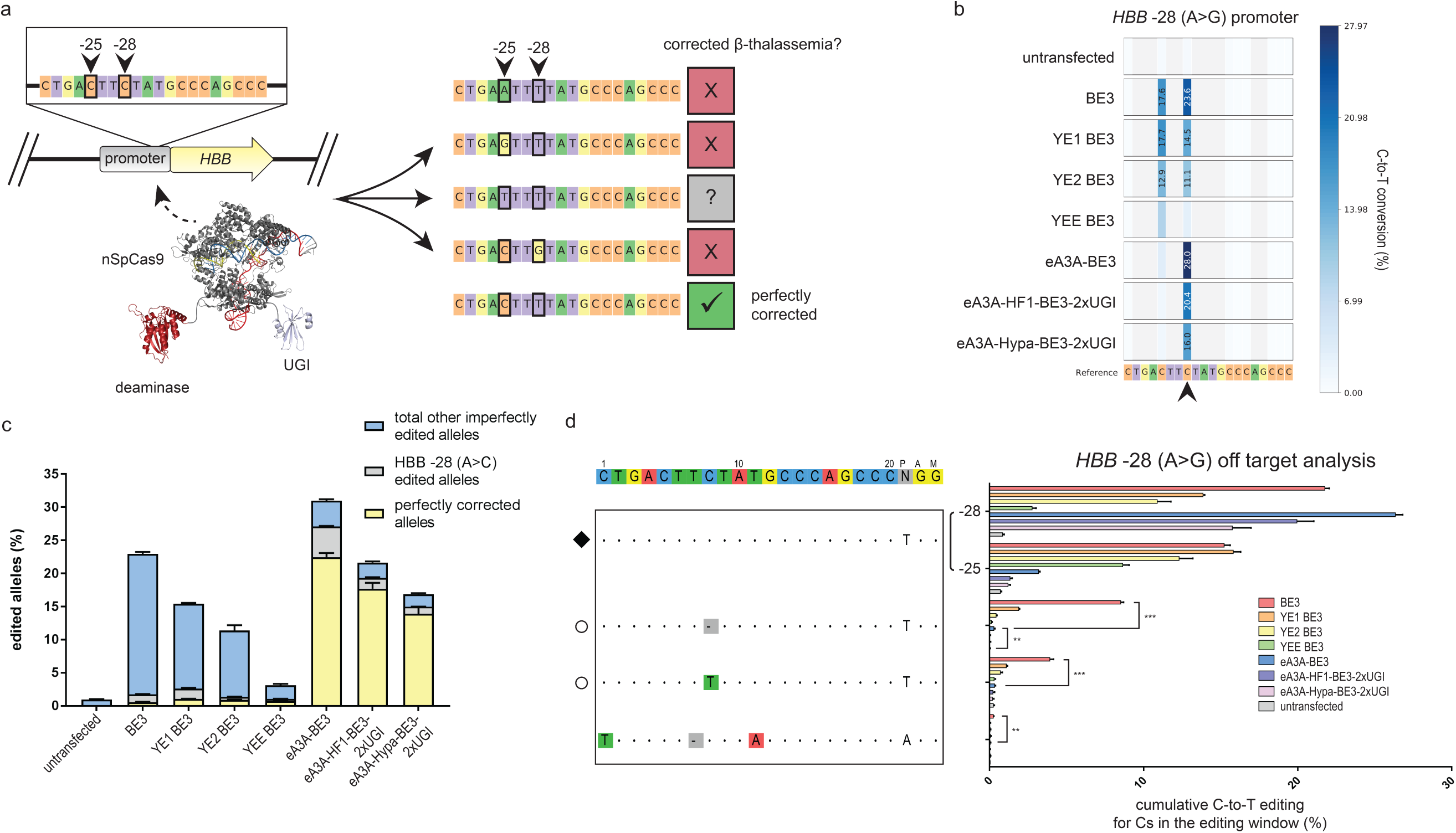
On- and off-target activities of eA3A-BE3 variants at a β-thalassemia-causing mutation *HBB* -28 (A>G) sequence in human cells. (A) Schematic of the *HBB* -28 (A>G) mutation and potential base editing outcomes when targeting Cs at -28 and -25 in the editing window of an *HBB*-targeting gRNA. Mutations to the bystander cytidine at the -25 position are deleterious and cause β-thalassemia phenotypes independent of the identity of the -28 nucleotide. (b) Heat maps showing C-to-T editing efficiencies for BE3, YE BE3s, and various A3A-BE3 variants at the *HBB* -28 (A>G) target site in an integrated reporter in human HEK293T cells. The -28 C is indicated with a black arrow. Editing efficiencies shown represent the mean of three biological replicates. (c) Graph showing the frequencies of perfectly corrected (-28 C to T only) and other imperfectly edited (-28 C to G or other edited Cs) alleles by BE3, YE BE3 variants, and eA3A-BE3 variants. (d) On- and off-target editing frequencies of the *HBB*-targeted gRNA with BE3, YE BE3 variants, or eA3A-BE3 variants. Percentage edits represent the sum of all edited Cs in the editing window and represent the mean of three biological replicates with error bars representing SEMs. Intended target sequence is shown at the top. On-target site is marked with a black diamond to the left and mismatches or bulges in the various off-target sites are shown with colored boxes or a dash in gray boxes, respectively. Off-target sites that lose the cognate TC motif within the editing window and thus might be expected to show lower off-target editing by eA3A, are noted with empty circles to the left. Asterisks indicate statistically significant differences in editing efficiencies observed between BE3 and eA3A-BE3 and between eA3A-BE3 and the untransfected control (* p < 0.05,** p < 0.005, *** p < 0.0005).

Our study provides an important proof-of-principle for how changing and engineering the cytidine deaminase in base editors can be used to optimize on-target precision and reduce off-target effects. We envision that a large suite of base editor fusions can be engineered by exploiting both the rich diversity of naturally occurring cytidine deaminase domains and the ability to modify the function and activity of these enzymes using protein engineering and evolution. In our study, mutation of the N57 residue in the human A3A deaminase was critical to restoring its native target sequence precision in the context of a base editor and also to lowering its off-target base editing activity. Introduction of additional mutations at A3A residues I96 and A71 further refined this precision, albeit at the expense of the desired cognate activity. Furthermore, the eA3A deaminase we engineered might be incorporated into and used to reduce the off-target effects of other base editor architectures that use different Cas9 orthologues for which high-fidelity variants have not yet been described (e.g., SaCas9 from *Staphylococcus aureus*^26^).

Relative to previously published studies, our strategy of using alternative and engineered cytidine deaminases provides a different and orthogonal approach to improve the precision of on-target editing. An earlier study introduced mutations into the rAPOBEC1 part of BE3 that shorten the editing window but this strategy narrows targeting range and does not permit predictable discrimination of base deamination when multiple cytidines are present in the window (as is the case with the β-thalassemia *HBB* -28 promoter mutation we successfully modified with eA3A-BE3 in this study). We also note that the YE BE3 variants that show the highest discrimination among multiple cytidines also typically show the greatest reductions in their overall base editing activity. One limitation of eA3A-BE3 is a decreased targeting range due to the increased sequence requirements flanking the target cytidine, a restriction that might be addressed by using engineered SpCas9 PAM recognition variants and naturally occurring Cas9 orthologues with different PAM specificities. In this regard, we have constructed an eA3A-BE3 derivative that uses an engineered SpCas9 variant that recognizes an NGA PAM^19^ and found that this fusion can efficiently edit sites bearing this alternative PAM (**Supplementary Fig. 8**). In addition, it may be possible to engineer or evolve different sequence specificities into APOBEC enzymes in the context of a base editor architecture, as has been done with APOBEC enzymes in isolation^13^,^27^. Thus, in the longer-term, we envision that the targeting range restriction might eventually be completely overcome by creating a larger series of different base editors that collectively recognize cytidines embedded in any sequence context.

## Online Methods

### Plasmids and oligonucleotides

Sequences of proteins and their expression plasmids used in this study are listed in the Supplementary Information. gRNA target sites and sequences of oligonucleotides used for on-target PCR amplicons for high throughput sequencing in this study can be found in **Supplementary Table 3**. Sequences of oligonucleotides used to investigate off-target editing sites can be found in **Supplementary Table 1**. BE expression plasmids containing amino acid substitutions were generated by PCR and standard molecular cloning methods. gRNA expression plasmids were constructed by ligating annealed oligonucleotide duplexes into MLM3636 cut with BsmBI. All gRNAs except those targeting the *HBB* -28 (A>G) and *CTNNB1* sites were designed to target sites containing a 5′ guanine nucleotide.

### Human cell culture and transfection

U2OS.EGFP cells containing a single stably integrated copy of the EGFP-PEST reporter gene and HEK293T cells were cultured in DMEM supplemented with 10% heat-inactivated fetal bovine serum, 2⏢mM GlutaMax, penicillin and streptomycin at 37⏢°C with 5% CO_2_. The media for U2OS.EGFP cells was supplemented with 400⏢μg⏢ml^-1^ Geneticin. Cell line identity was validated by STR profiling (ATCC), and cells were tested regularly for mycoplasma contamination. U2OS.EGFP cells were transfected with 750 ng of plasmid expressing BE and 250 ng of plasmid expressing sgRNA according to the manufacturer‘s recommendations using the DN-100 program and SE cell line kit on a Lonza 4-D Nucleofector. For HEK293T transfections, 75,000 cells were seeded in 24-well plates and 18 hours later were transfected with 600 ng of plasmid expressing BE and 200 ng of plasmid expressing sgRNA using TransIT-293 (Mirus) according to the manufacturer‘s recommendations. For all targeted amplicon sequencing and GUIDE-seq experiments, genomic DNA was extracted 72 h post-transfection. Cells were lysed in lysis buffer containing 100 mM Tris-HCl pH 8.0, 150 mM NaCl, 5 mM EDTA, and 0.05% SDS and incubated overnight at 55C in an incubator shaking at 250 rpm. Genomic DNA was extracted from lysed cells using carboxyl-modified Sera-Mag Magnetic Speed-beads resuspended in 2.5 M NaCl and 18% PEG-6000 (magnetic beads).

The HEK293T.HBB cell line was constructed by cloning a 200 base pair fragment of the *HBB* promoter upstream of an EF1a promoter driving expression of the puromycin resistance gene in a lentiviral vector. The *HBB* -28 (A>G) mutation was inserted by PCR and standard molecular cloning methods. The lentiviral vector was transfected into 293FS cells and media containing viral particles was harvested after 72 hours. Media containing viral particles was serially diluted and added to 10 cm plates with approximately 10 million HEK293T cells. After 48 hours, media was supplemented with 2.5 µg ml^-1^ puromycin and cells were harvested from the 10 cm plate with the fewest surviving colonies to ensure single copy integration.

### Off-target site selection and amplicon design

Two of the sites characterized here, *EMX1* site 1 and *FANCF*, were previously characterized by modified Digenome-seq, an unbiased approach to discover BE3-specific off-target sites. All off-targets discovered by modified Digenome-seq were investigated, and these sites represent the most comprehensive off-target characterization because they were discovered *de novo* using BE3. The *VEGFA* site 2 target is a promiscuous, homopolymeric gRNA that was previously characterized by GUIDE-seq. Because the *VEGFA* site 2 gRNA has over one hundred nuclease off-target sites, we selected the 20 off-target sites with the highest number of GUIDE-seq reads that also reside in loci for which we were able to design unique PCR amplification primers for characterization here. The *CTNNB1* and *HBB* -28 (A>G) gRNAs had not been previously characterized with respect to BE or nuclease off-target sites. We performed GUIDE-seq as previously described^17^ using these gRNAs to determine the SpCas9 nuclease off-target sites, and used Cas-OFFinder to predict all of the potential off-target sites with one RNA bulge and one mismatch. (GUIDE-seq and Cas-OFFinder analyses were performed using the hg38 reference genome.) This class of off-targets is more prevalent in BE3 relative to nucleases^16^, and thus sites that we were unlikely to discover by GUIDE-seq. Primers were designed to amplify all off-target sites such that potential edited cytidines were within the first 100 base pairs of Illumina HTS reads. A total of six primer pairs encompassing *EMX1* site 1, *VEGFA* site 2 and *CTNNB1* site 1 off-target sites did not amplify their intended amplicon and were thus excluded from further analysis.

### Statistical testing

All statistical testing was performed using two-tailed Student‘s t-test according to the method of Benjamini, Krieger, and Yekutieli without assuming equal variances between samples.

### Targeted amplicon sequencing

On- and off-target sites were amplified from ∼100 ng genomic DNA from three biological replicates for each condition. PCR amplification was performed with Phusion High Fidelity DNA Polymerase (NEB) using the primers listed in **Supplementary Tables 1 and 3**. 50 µl PCR reactions were purified with 1x volume magnetic beads. Amplification fidelity was verified by capillary electrophoresis on a Qiaxcel instrument. Amplicons with orthogonal sequences were pooled for each triplicate transfection and Illumina flow cell-compatible adapters were added using the NEBNext Ultra II DNA Library Prep kit according to manufacturer instructions. Illumina i5 and i7 indices were added by an additional 10 cycles of PCR with Q5 High Fidelity DNA Polymerase using primers from NEBNext Multiplex Oligos for Illumina (Dual Index Primers Set 1) and purified using 0.7x volume magnetic beads. Final amplicon libraries containing Illumina-compatible adapters and indices were quantified by droplet digital PCR and sequenced with 150 bp paired end reads on an Illumina MiSeq instrument. Sequencing reads were de-multiplexed by MiSeq Reporter then analyzed for base frequency at each position by a modified version of CRISPResso^28^. Indels were quantified in a 10 base pair window surrounding the expected cut site for each sgRNA.

### Expression of *HBB* -28 (A>G) gRNAs

In order to use eA3A BEs with the HF1 or Hypa mutations that decrease genome-wide off-target editing, it was necessary to use 20 nucleotides of spacer sequence in the gRNA with no mismatches between the spacer and target site^19^,^29^,^30^. We expressed the *HBB* -28 (A>G) gRNA from a plasmid using the U6 promoter, which preferentially initiates transcription at a guanine nucleotide at the +1 position. To preserve perfect matching between the spacer and target site, we appended a self-cleaving 5‘ hammerhead ribozyme that is able to remove the mismatched guanine at the 5‘ of the spacer^30^. This strategy rescued activity of HF1 eA3A BE3.9 or Hypa eA3A BE3.9 by approximately 1.4-fold compared to the gRNA with a 5‘ mismatched guanine (**Supplementary Fig. 9**).

## Data Availability

High–throughput sequencing reads will be deposited in the NCBI Sequence Read Archive database prior to publication.

## ACKNOWLEDGMENTS

This work was supported by grants from the National Institutes of Health (R35 GM118158 and RM1HG009490) and by the Desmond and Ann Heathwood MGH Research Scholar Award. J.G. was supported by the National Science Foundation Graduate Research Fellowship Program. We thank Alexander Sousa for advice on performing GUIDE-seq experiments, Peter Cabeceiras for assistance producing lentivirus, and James Angstman, Vikram Pattanayak, and Alexandra Mattei for helpful discussions and comments.

## AUTHOR CONTRIBUTIONS

J.M.G. conceived of the project, designed experiments and performed data analysis. O.C. performed all experiments with assistance from J.M.G. Data analysis was performed by M.K.C. with assistance from L.P. J.K.J. conceived of experiments and directed the research. J.K.J. and J.M.G. wrote the manuscript with input from all the authors.

## SUPPLEMENTARY TABLE LEGENDS

**Supplementary Table 1:** All off-target sites investigated by high-throughput sequencing in this study identified by name and sequence. Amplicon oligonucleotides represent the forward and reverse primers used to amplify each site from genomic DNA. Sites are organized by gRNA then by the method used to discover each off-target site.

**Supplementary Table 2:** Table of C to T editing efficiencies for each off-target site investigated in this study for BE3 or untransfected control. Statistically significant differences in editing efficiencies observed between BE3 and the untransfected control are indicated by p-value (p < 0.05).

**Supplementary Table 3:**. All on-target sites investigated by high-throughput sequencing in this study identified by name and sequence. Amplicon oligonucleotides represent the forward and reverse primers used to amplify each site from genomic DNA.

